# Effects of age on goal-dependent modulation of episodic memory retrieval

**DOI:** 10.1101/2020.09.04.283721

**Authors:** Sabina Srokova, Paul F. Hill, Rachael L. Elward, Michael D. Rugg

## Abstract

Retrieval gating refers to the ability to modulate the retrieval of features of a single memory episode according to behavioral goals. Recent findings demonstrate that younger adults engage retrieval gating by attenuating the representation of task-irrelevant features of an episode. Here, we examine whether retrieval gating varies with age. Younger and older adults incidentally encoded words superimposed over scenes or scrambled backgrounds that were displayed in one of three spatial locations. Participants subsequently underwent fMRI as they completed two memory tasks: the background task, which tested memory for the word’s background, and the location task, testing memory for the word’s location. Employing univariate and multivariate approaches, we demonstrated that younger, but not older adults, exhibited attenuated reinstatement of scene information when it was goal-irrelevant (during the location task). Additionally, in younger adults only, the strength of scene reinstatement in the parahippocampal place area during the background task was related to item and source memory performance. Together, these findings point to an age-related decline in the ability to engage retrieval gating.

## 1. Introduction

Episodic memory decline is well recognized as a prominent feature of cognitive aging (Grady et al., 2012; Nilsson, 2003; Nyberg et al., 2012). While one established factor driving this decline is reduced efficacy of encoding operations (e.g. Craik & Rose, 2012; Friedman & Johnson, 2014; Old & Naveh-Benjamin, 2008), the contribution of age differences in retrieval processing is less clear. A recently identified aspect of retrieval processing, termed ‘retrieval gating’, concerns the ability to regulate retrieved features belonging to a single memory episode according to their relevance to behavioral goals (Elward & Rugg, 2015). Retrieval gating refers to the finding that goal-relevant mnemonic features of an episode can be selectively reinstated, while reinstatement of irrelevant information is attenuated. This process is yet to be examined from the perspective of cognitive aging, raising the possibility that age-related episodic memory decline may, in part, be explained by inefficient gating of goal-irrelevant features of an episode.

Successful episodic memory retrieval requires the establishment of high-fidelity representations of episodes relevant to the retrieval goal. It has been proposed that retrieval of such episodes depends on strategic processes that bias the processing of retrieval cues to align it with the retrieval goal (Jacoby et al., 2005; Rugg, 2004; Rugg & Wilding, 2000). As such, memory search is optimized when cue processing can target goal-relevant episodes while avoiding retrieval of irrelevant information (memory ‘filtering’, in the terminology of Halamish et al. 2012). Indeed, recent findings point to the existence of mnemonic processes which work to attenuate the representation of goal-irrelevant associates of a retrieval cue in favor of relevant associates at the time of retrieval (Halamish et al., 2012; Wimber et al., 2015).

It has recently been demonstrated that, in addition to the processes discussed above that facilitate the retrieval and representation of goal-relevant episodes at the expense of irrelevant episodes, young adults appear also to be capable of retrieving a sub-set of the mnemonic features belonging to a *single* episode (Elward & Rugg, 2015; but see Kuhl et al., 2013 for conflicting findings). Elward & Rugg (2015) employed a paradigm akin to that of the present study in which participants incidentally encoded words superimposed over either scenes or a gray background. Each word-image pair was presented in one of three possible locations. At retrieval, participants underwent fMRI as they attempted to retrieve either the background or the location of studied test words. Goal-related modulation of episodic retrieval was investigated by examining task differences in the cortical reinstatement of scene information. Cortical reinstatement is a phenomenon characterized by the retrieval-related reactivation of neural patterns elicited during encoding (for reviews see Danker & Anderson, 2010; Rissman & Wagner, 2012; Rugg et al., 2015; Xue, 2018). In Elward and Rugg (2015), scene reinstatement effects were evident in parahippocampal and retrosplenial cortex - canonical scene-selective cortical regions (Aminoff et al. 2013; Bar 2004; Epstein & Baker, 2019) - when the retrieval task required a judgment about the nature of the background paired with the test words. Crucially, however, the effects were attenuated when scene information was irrelevant to the task, and instead, the task required a judgment about the test word’s study location. In light of prior research indicating that the strength of cortical reinstatement covaries with the amount and fidelity of recollected content (e.g. Thakral et al., 2015), these findings were interpreted as evidence that the retrieval of scene information was in some sense ‘gated’ when it was irrelevant to the retrieval goal.

Given the under-researched nature of retrieval gating and its underlying mechanisms, the term ‘gating’ as used here is not intended to imply a specific mechanism. Notably, we are agnostic as to whether gating is reflective of a biased memory search or a top-down control process which operates post-retrieval to attenuate the representation of goal-irrelevant mnemonic features. That said, if it is assumed that gating reflects an active control mechanism, the wealth of evidence demonstrating age deficits in top-down control motivates the hypothesis that older adults should have difficulty employing retrieval gating to regulate mnemonic content. For example, findings from prior behavioral studies examining age-differences in working memory (WM) suggest that the control processes that downregulate the representation of task-irrelevant information in WM are vulnerable to increasing age, consistent with the *inhibitory deficit hypothesis* of aging (Hasher & Zacks, 1988; Hasher et al., 1991; Lustig et al., 2001; Lustig et al., 2007, see also Campbell et al., 2020 and related articles in the same issue). However, despite the wealth of behavioral studies examining age deficits in WM, neuroimaging evidence for the inhibitory deficit hypothesis is relatively sparse. Nonetheless, extant findings suggest that older adults demonstrate reduced ability to strategically downregulate cortical activity in regions selectively responsive to specific classes of perceptual information (Chadick & Gazzaley, 2011; Chadick et al., 2014; Gazzaley et al., 2005, 2008; Weeks et al., 2020). For example, Chadick et al. (2014) reported that when participants were presented with overlapping images of a face and a scene in a delayed match to sample task, younger adults demonstrated attenuated activity in the parahippocampal cortex (relative to a ‘no-task’ baseline) when the scenes were task-irrelevant, and enhanced activity in the same region when they were task-relevant. In contrast, older adults did not demonstrate attenuated parahippocampal activity when the scenes were irrelevant, despite demonstrating enhancement to the same extent as young participants when the scenes were task-relevant.

In the present study, we examined whether, as might be anticipated on the basis of the foregoing brief review, retrieval gating becomes less efficient with increasing age. Younger and older adults undertook an incidental encoding task in which they elaboratively encoded words superimposed over scenes or scrambled backgrounds presented in one of three possible locations (cf. Elward and Rugg, 2015). Participants subsequently underwent fMRI as they completed two different retrieval tasks, requiring memory for either the background or the location of studied words. We expected to replicate the findings of Elward & Rugg (2015) that younger adults demonstrate attenuated scene reinstatement effects when scene information is not behaviorally relevant to the task. Crucially, we predicted that, relative to younger adults, older adults would be less able to modulate scene-related cortical reinstatement in accordance with the retrieval goal.

## 2. Materials and Methods

### 2.1. Participants

Twenty-seven younger and 30 older adults were recruited from communities surrounding The University of Texas at Dallas and were compensated $30/hour. All participants were right-handed, had normal or corrected-to-normal vision, and were fluent English speakers before the age of five. None of the participants had a history of cardiovascular or neurological disease, substance abuse, or diabetes, and none were using medication affecting the central nervous system at the time of participation. Potential participants were excluded from participation if they demonstrated evidence of cognitive impairment based on their performance on a neuropsychological test battery (see 2.2. Neuropsychological Testing).

Five younger and 6 older adults were excluded from the study and thus all subsequent fMRI analyses. Two younger adults and one older adult did not complete the scanning session due to claustrophobia or discomfort, and one younger adult was excluded due to technical difficulties during MRI scanning. Additionally, two younger and four older adults were excluded due to at-chance source memory performance in both the background and location tasks (see 2.4.2. Behavioral Data Analysis). Lastly, one older adult was excluded due to an incidental MRI finding. The final sample consisted of 20 younger adults (12 female, age range = 18 - 30 years) and 24 older adults (12 female, age range = 65 – 76 years).

### 2.2. Neuropsychological Testing

All participants completed a neuropsychological test battery on a separate day prior to participation in the fMRI session. The test battery consisted of the Mini-Mental State Examination (MMSE), The California Verbal Learning Test-II (CVLT; Delis et al., 2000), Wechsler Logical Memory (Test 1 and 2; Wechsler, 2009), the Symbol Digit Modalities test (SDMT; Smith, 1982), the Trail Making (Test A and B; Reitan and Wolfson, 1985), the F-A-S subtest of the Neurosensory Center Comprehensive Evaluation for Aphasia (Spreen and Benton, 1977), the Wechsler Adult Intelligence Scale – Revised (Forward and Backward digit span subtests; Wechsler, 1981), Category Fluency test (Benton, 1968), Raven’s Progressive Matrices (List 1; Raven et al., 2000), and a test of visual acuity. In addition, participants completed the Wechsler Test of Adult Reading (WTAR, Wechsler, 2001) or its revised version, the Wechsler Test of Premorbid Functioning (TOPF; Wechsler, 2011). Participants were excluded prior to the fMRI session if they performed > 1.5 SD below age norms on two or more non-memory tests, if they performed > 1.5 SD below at least one memory-based neuropsychological test, or if their MMSE score was < 27.

### 2.3. Experimental Procedure

#### 2.3.1. Materials

All experimental stimuli were presented using Cogent 2000 software (www.vislab.ucl.ac.uk/cogent_2000.php) implemented in Matlab 2012b (www.mathworks.com). The study phase was completed outside the scanner using a laptop and a button box identical to that used in the scanned memory task. During the MRI session, stimuli were projected on a translucent screen situated at the rear end of the scanner bore and viewed through a mirror placed on the head coil. The critical experimental stimuli comprised 240 concrete nouns, 120 colored images of urban and rural scenes (60 each), and 60 scrambled backgrounds created by randomly shuffling the pixels of 60 of the scene images. An additional 110 scenes, 110 scrambled backgrounds, and 110 object images were employed in a functional localizer task which was completed after the memory test. All images were scaled to 256 x 256 pixels. In addition to the critical stimuli, 39 nouns, 12 scenes, and 6 scrambled backgrounds were used either as practice stimuli or as filler trials at the beginning of a study or a test block. During the scanned test phase, the critical stimuli were randomly interspersed with 80 null trials, during which a black fixation cross was presented at the center of the display. Stimuli during the study and test phase were selected randomly to create 24 different stimulus sets, twenty of which were yoked between pairs of younger and older adults. Stimulus sets for the test phase were pseudo-randomized such that participants experienced no more than three consecutive new or old words studied against the same background or location, and no more than two consecutive null trials.

#### 2.3.2. Study Phase

At study, participants completed an incidental encoding task consisting of 180 trials divided across three study blocks of equal length, each taking 6 minutes 22 seconds to complete. Each block contained 60 words that were presented in one of three display locations (left, middle, right) and were superimposed over either a rural, an urban, or a scrambled background. Trials were equally divided such that a third of all words were presented over each background type and, independently, a third of the words were presented in each of the three locations. For example, across all study blocks, participants would study 60 words presented over an urban background, of which 20 would appear on the left side of the screen. A schematic of a study trial is presented in Figure 1-A. Each trial began with a black fixation cross presented for 200 ms in the square corresponding to the location in which the upcoming word-image pair was to be presented. The cross was replaced by the study word, followed 200 ms later by its background. The word and image pair remained together on the screen for 5500 ms, and participants were instructed to imagine a scenario in which the object denoted by the word is moving around or interacting with the background. Participants rated the vividness of this imagined scenario on a three-point scale by responding on the button box with the index, middle, and ring fingers (1 = not vivid, 2 = somewhat vivid, 3 = very vivid). The inter-trial-interval, during which only the three grey squares and response prompts remained on the screen, lasted 400 ms.

**Figure 1.**
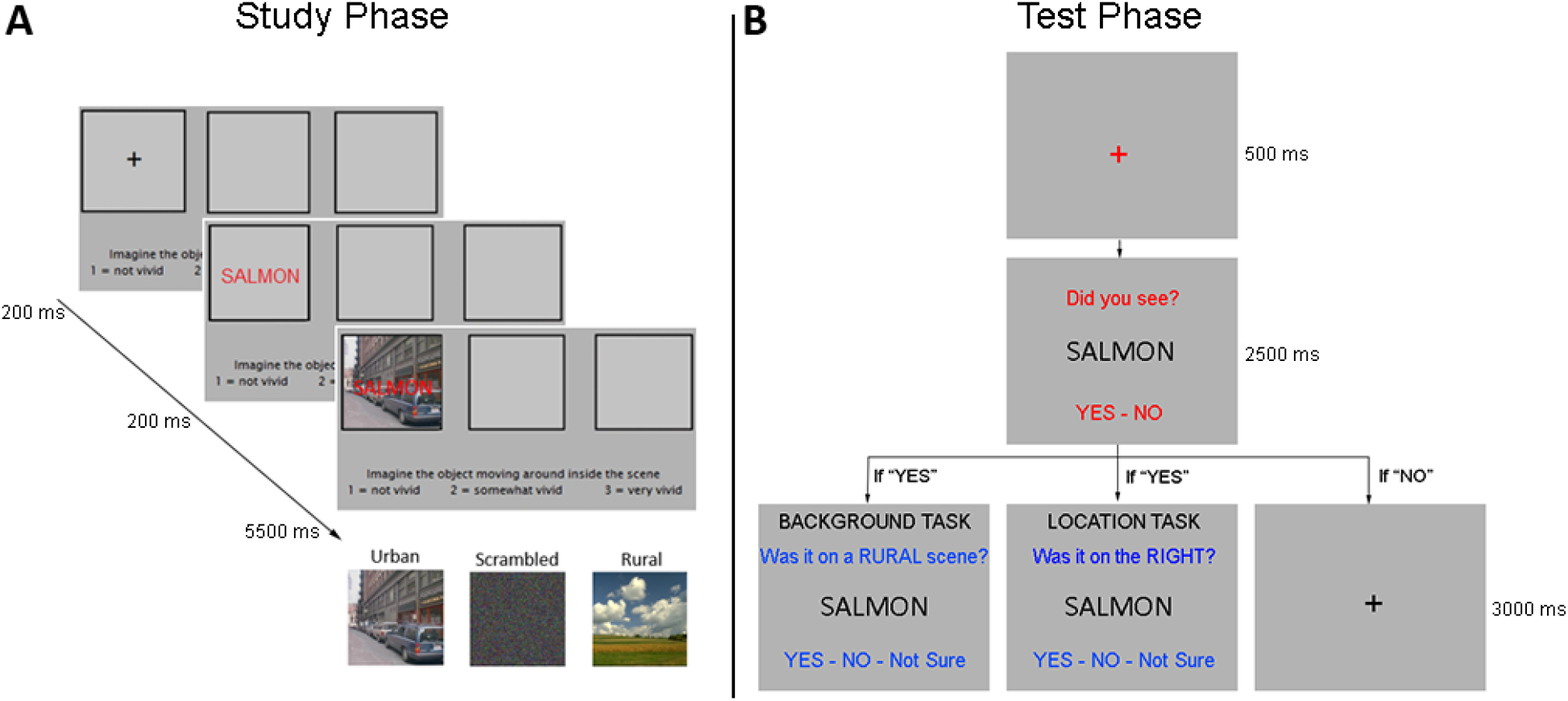
Task schematics for the Study (A) and Test Phases (B). The study phase was completed on a laptop while the memory test was completed inside an MRI scanner. The memory test consisted of two retrieval tasks: the background and the location task.

#### 2.3.3. Test Phase

Participants were trained on the memory test after completing the study task (hence, encoding was incidental). The test phase was completed inside the scanner approximately 30 minutes following the completion of the study task, and consisted of 240 critical trials (comprising 180 studied and 60 unstudied words) which were divided equally between 5 blocks of ‘background’ and 5 blocks of ‘location’ tests distributed across 5 scanning runs. Each scanning run lasted 8 minutes 15 seconds and contained a single location and a background task block. In each block, participants were presented with 24 critical test words (6 new, 18 old) intermixed with 8 null trials. The 18 old words were balanced such that a third of all old trials in a block were associated with each of the three background and location contexts at study. Figure 1-B illustrates a schematic of a single test trial and the response alternatives for the two retrieval tasks. Each test trial began with a red fixation cross presented for 500 ms in the middle of the screen, which was subsequently replaced by the test word for 2500 ms. At the word onset, the response prompt “Did you see?” appeared above the word, and cues “Yes – No” were presented underneath. Participants indicated whether they remembered seeing the word at study using the index and middle fingers of the right hand. For each word endorsed old (i.e. following a “Yes” response), participants were presented with a follow-up source memory prompt that was displayed for 3000 ms. For each word endorsed new (i.e. following a “No” response), a black fixation cross was displayed for 3000 ms until the end of the trial. The inter-trial-interval, during which the black fixation cross remained on the screen, lasted 1000 ms.

In the location task block, participants were required to recall the location of the studied word according to one of the following prompts presented above the word: “Was it on the LEFT?” or “Was it on the RIGHT?”. In the background task block, participants recalled the background scene following either “Was it on an URBAN scene?” or “Was it on a RURAL scene?”. The response cues “Yes – No – Not Sure” were presented below the word, and participants made responses with their right hand using the index, middle, and ring fingers. Trials requiring a correct “Yes” source response are henceforth termed ‘target’ trials, whereas trials requiring a correct “No” response are termed ‘non-target’ trials. The target trial type was counterbalanced across participants such that the target location and target background type remained constant across all trials. Thus, if a participant’s location target type corresponded with trials studied on the right side on the screen, they would be presented with the prompt “Was it on the RIGHT?” on each location task trial. Similarly, for participants whose target background trials comprised words studied over the rural scenes, the prompt “Was it on a RURAL scene?” would be shown across all background task trials. The mapping of the responses to fingers was counterbalanced across participants with the constraint that the “Not Sure” response was mapped onto the ring or index finger. If the “Not Sure” response was assigned to the ring finger, the “Yes” and “No” responses were assigned to the index and middle fingers, respectively. Otherwise, the “Yes” and “No” responses were assigned to the middle and ring fingers, respectively. The order of the response cues displayed on the screen was adjusted accordingly. Lastly, the ordering of the retrieval tasks was counterbalanced, such that first half of each scanning run corresponded with the location task for half of the participants, and the background task for the remainder. Participants were informed of the task they were about to complete by a reminder that was displayed prior to the onset of each task block.

#### 2.3.4. Functional Localizer

Following the completion of the test phase and the acquisition of a structural MRI scan, participants completed the functional localizer task. The task comprised 5 scanner runs. Each run lasted 2 minutes 51 seconds and consisted of 6 blocks of 11 trial-unique images that comprised either scenes, scrambled backgrounds, or common objects. A single image of a face was interspersed randomly within each block. Participants were instructed to press a button whenever they saw the face image. Each image was presented for 750 ms and followed by a black fixation cross lasting for 250 ms. The blocks of images were separated by a 12 second interval during which a black fixation cross was continuously present in the middle of the display.

### 2.4. Data Acquisition and Analysis

#### 2.4.1. Statistical Analysis

All statistical analyses and data visualization were performed using R software (R Core Team, 2020). Analyses of variance were performed with the package afex (Singmann et al., 2016) with the degrees of freedom and p-values corrected for nonsphericity with the Greenhouse and Geisser (1959) procedure. Correlations were performed with the cor.test function, linear regressions were performed with the lm function, and pairwise comparisons were performed with the function t.test, all in base R. All t-tests were two-tailed except for the one-sample t-tests that tested whether reinstatement and similarity indices were significantly greater than zero (i.e. to determine whether reinstatement effects were reliable). Effect sizes for the analyses of variance are reported as partial-*η*^2^ and effect sizes for t-tests are reported as Cohen’s d. Figures plotting fMRI data were created using the package ggplot2 (Wickham, 2016). All tests were considered significant at p < 0.05.

#### 2.4.2. Behavioral Data Analysis

Item memory performance was evaluated by examining recognition memory performance with respect to the “Did you see?” prompt. Item recognition (Pr) was computed as the difference between the proportion of correctly recognized old words (item hits) and the proportion of new trials which were incorrectly endorsed as old words (false alarms):

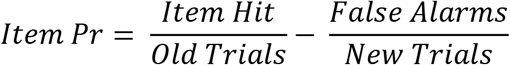

Source memory performance (pSR; probability of source recollection) was evaluated in terms of correctly judging whether or not a word endorsed old had been studied in association with the target background or location. pSR was estimated with a modified single high-threshold model (Snodgrass and Corwin, 1988; see also Gottlieb et al., 2010; Mattson et al., 2014) using the following formula:

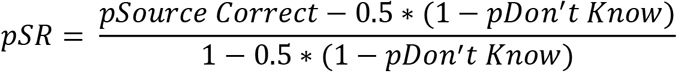

Here, ‘pSource Correct’ and ‘pDon’t know’ refer to the proportion of correctly recognized old trials which received an accurate source memory judgement or a ‘Not Sure’ response, respectively.

#### 2.4.3 MRI Data Acquisition and Preprocessing

Functional and structural MRI data were acquired using a 3T Philips Achieva MRI scanner (Philips Medical Systems, Andover, MA) equipped with a 32-channel head coil. Functional images during the test phase were acquired with a T2*-weighted, blood-oxygen-level-dependent echoplanar imaging (EPI) sequence with a multiband factor of two (flip angle = 70°, field of view [FOV] = 200 x 240 mm, repetition time [TR] = 1.6 s, echo time [TE] = 30 ms). EPI volumes consisted of 44 slices at a voxel size of 2.5 x 2.5 x 2.5 mm with a 0.5 mm interslice gap. The slices were acquired in an interleaved order and oriented parallel to the anterior-posterior commissure line. The protocol for the functional localizer was identical to the test phase protocol except for the repetition time (TR = 1.5 s). Structural images were acquired with a T1-weighted MPRAGE sequence (FOV = 256 x 256 mm, 1 x 1 x 1 mm isotropic voxels, sagittal acquisition).

The MRI data were preprocessed and analyzed using Statistical Parametric Mapping (SPM12, Wellcome Department of Cognitive Neurology, London, UK) and custom Matlab code. The functional data were realigned to the mean EPI image and slice-time corrected using *sinc* interpolation with reference to the 12^th^ acquired slice. The images were then normalized to a sample-specific template according to previously published procedures to ensure an unbiased contribution of each age group to the template (de Chastelaine et al. 2011, 2016). Prior to region-of-interest (ROI) definition, the time series of each localizer run were concatenated using the *spm_fmri_concatenate* function and smoothed with a 6 mm full-width half-maximum Gaussian kernel. The reinstatement indices extracted from the test-phase data, as well as the test and localizer time series used in the pattern similarity analysis, were derived from unsmoothed data.

#### 2.4.4 MRI Data Analysis

##### 2.4.4.1 ROI Selection

The data from the localizer task were analyzed in two stages prior to defining the ROIs. First, a separate 1^st^-level GLM was constructed for each participant by modeling 3 events of interest: scene blocks, object blocks, scrambled blocks. Each block was modeled with a 12s duration boxcar regressor convolved with a canonical hemodynamic response function (HRF) with temporal and dispersion derivatives, onsetting concurrently with the presentation of the first stimulus in the block. Covariates of no interest comprised 6 regressors reflecting motion-related variance (rigid-body translation and rigid-body rotation) and the mean signal of each session run. Additionally, motion spikes with translational displacement of > 1 mm or rotational displacement of > 1° were modeled as additional covariates of no interest. The subject-level parameter estimates were carried over to a 2^nd^-level GLM that took the form of a 2 (age group: younger, older) x 3 (stimulus type: scene, object, scrambled) mixed effects ANOVA model. Age group was included in the ANOVA to ascertain that the age group-by-stimulus type interaction did not reveal additional scene-selective clusters outside of those described below.

The ROIs were derived using the conjunction of scene > object and scene > scrambled contrasts at the 2^nd^-level, both thresholded at p < 0.0005 (uncorrected), 50 voxels. [Note that when the ROIs were defined using either a stricter or a more liberal threshold (p < 0.0001 and p < 0.01 respectively) the results reported in the fMRI analyses below were unchanged]. The contrasts employed the simple effects of stimulus type to ensure that the ROIs were unbiased with respect to age group. This procedure identified scene-selective clusters in the parahippocampal place area (PPA) and retrosplenial cortex (RSC) bilaterally (Figure 2-A; peak MNI coordinates presented in Table 1). The left and right PPA were delimited by restricting the cluster with a combination of anatomical masks corresponding to the parahippocampal and fusiform gyri provided by the Neuromorphometrics atlas in SPM12. We created an RSC mask by searching the Neurosynth database for the term “retrosplenial” (search in August 2019, search results FDR-corrected at p < 0.00001; Yarkoni et al., 2011). The resulting mask was used to restrict the outcome of the localizer contrast, thereby generating the left and right RSC ROIs (Figure 2-B).

**Figure 2:**
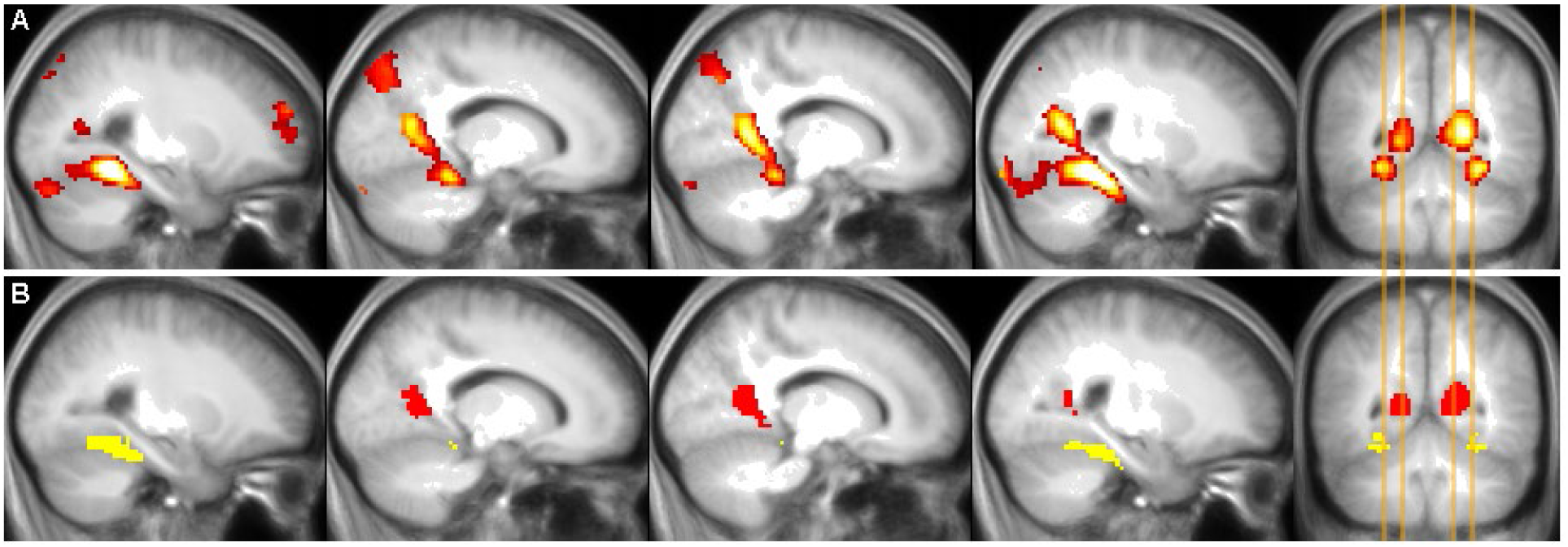
**A:** Functional localizer data illustrating scene-selective clusters used to define the ROIs employed in the fMRI data analysis. The illustrated clusters represent the conjunction of the scene > object and scene > scrambled background contrasts before masking (see main text). **B:** Scene-selective PPA and RSC ROIs derived by masking the clusters in 2-A (see main text).

**Table 1:** Cluster peak MNI coordinates and the number of voxels in of each ROI derived from the functional localizer. The peak MNI coordinates were obtained from the scene > object contrast which was inclusively masked with the scene > scrambled contrast.

|  | ROI Size<br>(voxels) | Peak MNI Coordinates |  |  |
| --- | --- | --- | --- | --- |
|  |  | X | Y | Z |
| R. Parahippocampal place area | 212 | 27 | -47 | -12.5 |
| L. Parahippocampal place area | 177 | -28 | -49.5 | -10 |
| R. Retrosplenial cortex | 200 | 19.5 | -57 | 10 |
| L. Retrosplenial cortex | 121 | -18 | -57 | 5 |

R = Right, L = Left

##### 2.4.4.2. Univariate Reinstatement Index

The unsmoothed functional data from the test phase were concatenated and subjected to a ‘least-squares-all’ GLM analysis (Mumford et al., 2014; Rissman et al., 2004) to estimate the BOLD response elicited on each test trial. Each trial was modeled as a separate event of interest with a delta function time-locked to stimulus onset and convolved with a canonical HRF. (The employment of the delta function follows the approach adopted by Elward and Rugg (2015). We note however that the findings were unchanged when the data were modeled with a 2.5s boxcar that tracked the duration of the ‘did you see’ prompt that onset concurrently with the test item). Covariates of no interest consisted of the aforementioned 6 motion regressors reflecting translational and rotational displacement, and the regressors of session-specific means with the first run as the intercept.

‘Reinstatement indices’ based on the resulting single-trial β-weights in each scene-selective ROI, separately for the trials of the two retrieval tasks, were computed in a manner akin to a previously described ‘differentiation index’ (Koen et al., 2019; Srokova et al., 2020; Voss et al., 2008; Zebrowitz et al., 2016). The reinstatement index operationalizes retrieval-related reinstatement of scene information in terms of an effect size by computing the difference between the mean BOLD response associated with test words studied over scene backgrounds versus words studied over scrambled backgrounds, and then dividing the difference by pooled standard deviation:

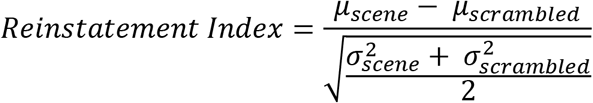

Reinstatement indices greater than zero reflect a greater mean BOLD response for words studied over scenes relative to words studied over scrambled backgrounds, and as such, are indicative of retrieval-related cortical reinstatement of scene information. Importantly, because of the scaling function, the reinstatement index is insensitive to individual differences in the gain of the HRF which mediates the relationship between neural activity and the resulting fMRI BOLD signal. Thus, the reinstatement index is unaffected by age-related differences in HRF gain (see, for example, Liu et al., 2013)

Following Elward and Rugg (2015), the test trials employed in the computation of the reinstatement index were those on which test words were correctly recognized as previously studied, irrespective of the accuracy of the subsequent source memory judgment. The inclusion of all correctly recognized trials allows for a fair comparison of the scene reinstatement effects in the two tasks (see Elward and Rugg, 2015). Trials on which test words had been paired with target scenes were however excluded from these fMRI analyses, such that the analyses were performed only for recognized test words associated with non-target backgrounds (i.e. non-target and scrambled scenes). This approach was adopted to reduce the influence of any confound arising from the fact that scrambled trials in the background task were always non-targets (and hence should always have received a ‘No’ source response), whereas scene trials were a mixture of non-targets (‘No’ response) and targets (‘Yes’ response). Thus, we aimed to ensure that the analysis of scene reinstatement in the background task was not confounded by the differential responses associated with test words associated with scrambled as opposed to target scenes. To ensure that the contrast between scene reinstatement effects across the two tasks was between scenes belonging to same sub-category (rural or urban), analysis of scene reinstatement effects in the location task was also restricted to test words associated with non-target scenes. To the extent that item recognition performance was equivalent across the two retrieval tasks, this approach also balanced the number of background and location task trials contributing to the computation of the respective reinstatement indices.

##### 2.4.4.3. Pattern Similarity Analysis

Pattern similarity analysis (PSA) was performed to complement the analyses of the univariate reinstatement index (cf. Koen et al., 2019; Srokova et al., 2020). Scene reinstatement was operationalized in terms of shared neural patterns between test trials and a scene-specific voxel-wise profile derived from the functional localizer. To this end, for each ROI, voxel-wise test-phase single-trial β-weights extracted from the least-squares-all GLM analysis described above were correlated with the voxel-wise β-weight profile derived from the scene > scrambled contrast conducted on the localizer data (although we note that the PSA results remain unchanged if the scene > scrambled + object contrast is employed instead). Other than being performed on unsmoothed data, the localizer data were modeled as described previously, and the scene > scrambled profiles were derived for each participant from the first-level GLMs. PSA was conducted on the same test trials as those employed for the analyses of reinstatement indices described in the previous section. Thus, only those trials containing correctly recognized test items associated with non-target backgrounds at study were included in these analyses. Scene-related cortical reinstatement was operationalized as the difference between the across-trial mean Fisher z-transformed correlation between the localizer contrast and non-target scene trials, and the mean Fisher z-transformed correlation between the localizer contrast and all scrambled trials (see Figure 3). Similarity scores were computed separately for the location and background tasks in each ROI. Similarity indices greater than zero are indicative of scene reinstatement at test. Importantly, as for the reinstatement index, the similarity index is insensitive to individual differences in HRF gain.

**Figure 3:**
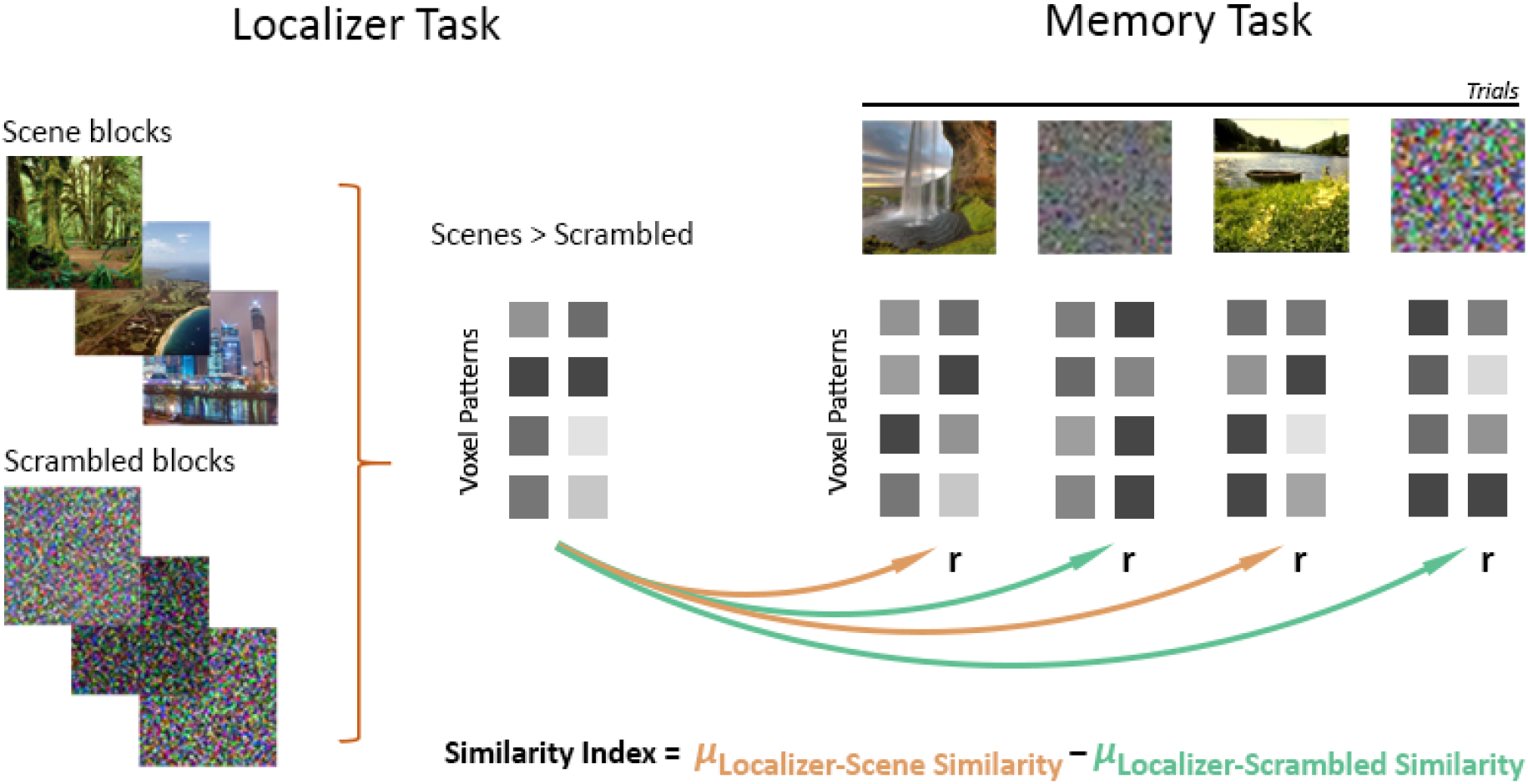
Schematic of the PSA. Similarity indices were computed separately for each task as the difference between the mean correlation between the localizer contrast and all non-target scene trials (localizer-scene similarity) and the mean correlation between the localizer and all scrambled trials (localizer-scrambled similarity).

In light of the literature describing age differences in neural specificity (so-called ‘age-related neural dedifferentiation’, for review see Koen & Rugg, 2019), we performed a control analysis to ensure that the PSA results presented below were not driven by age differences in scene selective activity identified by the functional localizer. To accomplish this, individual localizer blocks were modeled separately as 12s boxcars onsetting concurrently with the first stimulus of each block. A PSA was then performed on the resulting block-wise β-weights by computing a within – between similarity metric separately in each scene-selective ROI. The within similarity metric comprised the average Fisher z-transformed Pearson’s correlation between a given scene block and all other scene blocks. The between similarity metric was the average Fisher z-transformed correlation between a given scene block and all scrambled blocks. A given block was never correlated with another block belonging to the same scanner run to avoid potential bias arising from carry-over effects (Mumford et al., 2014). The within – between similarity indices were then entered into a 2 (age group) by 4 (ROI) mixed ANOVA. This revealed a main effect of ROI (F_(2.41, 98.93)_ = 45.244, p < 0.001, partial-η^2^ = 0.525), but neither the main effect of age nor the age-by-ROI interaction term reached significance (F_(1, 41)_ = 1.648, p = 0.206, partial-η^2^ = 0.039, and F_(2.41, 98.93)_ = 1.941, p = 0.140, partial-η^2^ = 0.045, respectively). These results indicate that scene-related neural specificity, as indexed by PSA, did not differ between younger and older adults in the localizer task. Thus, the localizer data provide an unbiased baseline for the examination of age differences in PSA metrics of scene reinstatement during the two retrieval tasks.

## 3. Results

### 3.1. Neuropsychological Test Results

Demographic data and performance on the neuropsychological test battery are reported in Table 2. Analysis of the test scores revealed that younger adults outperformed older adults on the Wechsler Logical Memory subtest I, achieved a greater overall number of recognition hits on CVLT, and demonstrated faster processing speed as reflected by their performance on Trails Subtest A and the SDMT. Younger adults also demonstrated better performance relative to older adults on Raven’s Progressive Matrices.

**Table 2.** Demographic data and performance on the neuropsychological test battery: Mean (SD) and age differences (significant age differences denoted by *).

|  | Younger Adults | Older Adults | Age difference (p) |
| --- | --- | --- | --- |
| N (male/female) | 8/12 | 12/12 |  |
| Age | 24.55 (3.64) | 68.50 (3.19) |  |
| Years of Education | 15.95 (1.64) | 17.25 (2.66) | 0.054 |
| MMSE | 29.05 (0.89) | 29.08 (0.88) | 0.902 |
| CVLT Short Delay - Free | 11.90 (2.45) | 10.29 (3.26) | 0.069 |
| CVLT Short Delay - Cued | 12.95 (2.06) | 11.63 (2.78) | 0.077 |
| CVLT Long Delay - Free | 12.70 (2.16) | 11.17 (3.13) | 0.062 |
| CVLT Long Delay - Cued | 12.60 (2.37) | 11.83 (2.57) | 0.309 |
| CVLT Recognition Hits | 15.50 (0.69) | 14.58 (1.38) | 0.007 * |
| CVLT False Alarms | 1.45 (1.64) | 2.13 (2.35) | 0.269 |
| F-A-S | 45.95 (11.01) | 46.33 (10.68) | 0.908 |
| WMS Logical Memory I | 31.55 (6.83) | 27.79 (4.60) | 0.044 * |
| WMS Logical Memory II | 29.50 (5.42) | 26.79 (4.63) | 0.086 |
| Digit-symbol Substitution Test | 63.65 (10.73) | 50.54 (7.59) | 0.001 * |
| Trails A (s) | 20.62 (7.50) | 29.46 (10.23) | 0.002 * |
| Trails B (s) | 51.85 (32.71) | 62.04 (20.29) | 0.235 |
| Digit Span<sup>1</sup> | 18.70 (2.56) | 18.38 (3.92) | 0.743 |
| Category Fluency Test | 26.00 (6.68) | 22.83 (4.07) | 0.074 |
| TOPF / WTAR<sup>2</sup> | 109.55 (10.09) | 113.58 (7.45) | 0.147 |
| Raven's Progressive Matrices I | 10.85 (0.93) | 9.63 (1.81) | 0.007 * |

<sup>1</sup>Digit span corresponds to the sum of forward and backward digit span.
<sup>2</sup> Presented as a standardized score computed from WTAR or TOPF raw scores

### 3.2. Behavioral Performance

Behavioral performance including vividness ratings at study, study and test reaction times (RTs), and memory performance are presented in Table 3. Vividness ratings and mean RTs at study were binned according the trials’ background context (scrambled backgrounds, target scenes, non-target scenes). Vividness data from one older adult were not recorded due to a technical malfunction. The vividness ratings and RTs were submitted to a 2 (age group) x 3 (background context) mixed factorial ANOVA. In the case of rated vividness, the main effects of age group, background context, and the two-way interaction were in each case not significant (age group: F_(1,41)_ = 0.296, p = 0.589, partial-η^2^ = 0.007; background context: F_(1.41, 57.72)_ = 1.978, p = 0.159, partial-η^2^ = 0.046; age-by-background interaction: F_(1.41, 57.72)_ = 0.596, p = 0.498, partial-η^2^ = 0.014). Analysis of the study RTs revealed a significant main effect of age group (F_(1,41)_ = 5.590, p = 0.023, partial-η^2^ = 0.120), reflective of faster RTs in the younger cohort. The main effect of background context was also significant (F_(1.23, 50.29)_ = 51.206, p < 0.001, partial-η^2^ = 0.555), reflecting faster RTs to scrambled trials relative to either scene type (scrambled vs. target scene: t_(42)_ = 8.102, p < 0.001, d = 1.236; scrambled vs. non-target scene: t_(42)_ = 6.857, p < 0.001, d = 1.046; target scene vs. non-target scene: t_(42)_ = 0.676, p = 0.503, d = 0.103). These background effects did not differ significantly between the age groups (two-way interaction: F_(1.23, 50.29)_ = 3.041, p = 0.079, partial-η^2^ = 0.069).

**Table 3.** Mean (SD) memory performance and RT at test.

|  | Younger Adults | Older Adults |
| --- | --- | --- |
| *Study Phase* |  |  |
| Vividness Ratings |  |  |
| Scrambled background | 2.32 (0.41) | 2.19 (0.56) |
| Target scene | 2.39 (0.31) | 2.37 (0.40) |
| Non-target scene | 2.34 (0.38) | 2.32 (0.43) |
| Study RTs (ms) |  |  |
| Scrambled background | 2704.08 (745.65) | 3012.18 (580.91) |
| Target scene | 3077.41 (734.65) | 3661.12 (616.52) |
| Non-target scene | 3101.65 (712.29) | 3603.46 (685.27) |

| *Test Phase* | *Location Task* | *Background Task* | *Location Task* | *Background Task* |
| --- | --- | --- | --- | --- |
| Item Hit Rate | 0.81 (0.12) | 0.80 (0.14) | 0.72 (0.12) | 0.71 (0.10) |
| False Alarm Rate | 0.15 (0.15) | 0.16 (0.12) | 0.14 (0.09) | 0.12 (0.08) |
| Item Recognition (Pr) | 0.66 (0.21) | 0.64 (0.18) | 0.58 (0.15) | 0.59 (0.15) |
| Source Memory (pSR) | 0.23 (0.16) | 0.49 (0.19) | 0.16 (0.13) | 0.38 (0.19) |
| Proportion Source Correct |  |  |  |  |
| Scrambled background | 0.52 (0.17) | 0.63 (0.21) | 0.47 (0.18) | 0.56 (0.25) |
| Target scene | 0.48 (0.15) | 0.70 (0.17) | 0.41 (0.15) | 0.65 (0.16) |
| Non-target scene | 0.49 (0.17) | 0.73 (0.17) | 0.39 (0.15) | 0.66 (0.19) |
| Proportion Don't Know |  |  |  |  |
| Scrambled background | 0.31 (0.18) | 0.25 (0.18) | 0.33 (0.22) | 0.18 (0.18) |
| Target scene | 0.28 (0.18) | 0.15 (0.12) | 0.33 (0.19) | 0.14 (0.13) |
| Non-target scene | 0.27 (0.17) | 0.16 (0.15) | 0.33 (0.22) | 0.14 (0.16) |
| Test RT (ms) |  |  |  |  |
| Item responses | 1298.09 (122.09) | 1300.49 (125.39) | 1506.76 (160.23) | 1479.31 (162.99) |
| Source responses | 995.97 (247.21) | 1055.38 (252.48) | 1439.47 (319.65) | 1383.78 (285.93) |

*Item recognition memory computed as the difference between hit and false alarm rates*
*Source memory computed using the single high-threshold model described in Behavioral Data Analysis*

Memory performance was analyzed with separate 2 (age group) x 2 (retrieval task) ANOVAs for item recognition (Item Pr) and source memory performance (pSR). With respect to item recognition, the main effects of age group and retrieval task were both non-significant (age group: F_(1,42)_ = 1.767, p = 0.191, partial-η^2^ = 0.040; retrieval task: F_(1,42)_ = 0.056, p = 0.814, partial-η^2^ = 0.001), and the age group-by-task interaction also failed to reach significance (F_(1,42)_ = 1.321, p = 0.257, partial-η^2^ = 0.030). The ANOVA of source accuracy scores revealed a significant main effect of age group (F_(1,42)_ = 5.458, p = 0.024, partial-η^2^ = 0.115), a main effect of retrieval task (F_(1,42)_ = 63.417, p < 0.001, partial-η^2^ = 0.602), but no age group-by-task interaction (F_(1,42)_ = 0.291, p = 0.593, partial-η^2^ = 0.007). The main effect of task was driven by superior accuracy in the background relative to the location task across both age groups. Additionally, older adults’ source accuracy was lower than that of their younger counterparts on both tasks.

To examine the effects of background context on memory performance (analogous to the analysis of the study data), the following analysis examined the proportion of correctly recognized items that went on to receive a correct source memory judgement (pSource correct) in test trials binned according to their background context at study (scrambled backgrounds, target scenes, non-target scenes). Effects of study context were examined with a 2 (age group) x 2 (retrieval task) x 3 (background context) mixed factorial ANOVA. This revealed a significant main effect of task (F_(1, 42)_ = 66.992, p < 0.001, partial-η^2^ = 0.615), but no main effect of age group (F_(1, 42)_ = 3.239, p = 0.079, partial-η^2^ = 0.072) or background context (F_(1.81, 75.87)_ = 0.693, p = 0.489, partial-η^2^ = 0.016). Additionally, the ANOVA revealed a significant task-by-background context interaction (F_(1.84, 77.16)_ = 10.061, p < 0.001, partial-η^2^ = 0.193), along with non-significant interactions between background context and age group (F_(1.81, 75.87)_ = 0.162, p = 0.830, partial-η^2^ = 0.004), age group-by-task (F_(1, 42)_ = 0.056, p = 0.814, partial-η^2^ = 0.001) and a non-significant three-way interaction (F_(1.84, 77.16)_ = 0.199, p = 0.802, partial-η^2^ = 0.005).

The significant task-by-background context interaction was examined with additional 2 (age group) x 3 (background context) ANOVAs performed separately for the location and background tasks. In the location task, the ANOVA identified a main effect of background context (F_(1.95, 81.79)_ = 3.983, p = 0.023, partial-η^2^ = 0.087), while the main effect of age group and the two-way interaction were not significant (F_(1, 42)_ = 2.861, p = 0.098, partial-η^2^ = 0.064; F_(1.95, 81.79)_ = 0.483, p = 0.614, partial-η^2^ = 0.011, respectively). The main effect of background context was driven by better location memory for words studied over scrambled backgrounds relative to either non-target (t_(43)_ = 2.869, p = 0.006, d = 0.433) or target scenes (t_(43)_ = 2.192, p = 0.034, d = 0.330). The corresponding ANOVA for the background task also identified a main effect of background context (F_(1.74, 72.98)_ = 5.534, p = 0.008, partial-η^2^ = 0.116). The main effect of age group and the two-way interaction were again not significant (F_(1, 42)_ = 1.749, p = 0.193, partial-η^2^ = 0.040; F_(1.74, 72.98)_ = 0.044, p = 0.939, partial-η^2^ = 0.001, respectively). In this case, the main effect of context was driven by lower source memory for words studied in association with scrambled backgrounds relative to non-target (t_(43)_ = 3.465, p = 0.001, d = 0.522) or target scenes (t_(43)_ = 2.210, p = 0.032, d = 0.333).

Two 2 (age group) x 2 (retrieval task) x 3 (background context) mixed ANOVAs were also employed to evaluate RTs for the item and source memory judgements. The ANOVA examining item recognition RTs identified a main effect of age group (F_(1,42)_ = 23.729, p < 0.001, partial-η^2^ = 0.361); the remaining main effects and the interactions were not significant (ps > 0.080). Thus, while older adults were slower overall in making their recognition judgements, neither background context nor retrieval task moderated RTs in either age group. The ANOVA for source memory RTs again revealed a main effect of age group (F_(1, 42)_ = 25.512, p < 0.001, partial-η^2^ = 0.378), but in this case the effect was modified by an age group-by-task interaction (F_(1, 42)_ = 5.517, p = 0.024, partial-η^2^ = 0.116). The main effects of background context and task, and the remaining two- and three-way interactions were not significant (ps > 0.165). These results indicated that, akin to item recognition, older adults were slower overall to make source memory judgements relative to younger adults. Additionally, the age group-by-task interaction for source RTs indicated that age differences in source RTs were sensitive to retrieval task. Follow-up comparisons demonstrated that the interaction was driven by faster source judgements in the location relative to the background task in younger adults (p = 0.027), whereas the task effect was not significant in the older adults (p = 0.173). Consistent with item recognition, the source memory RTs were insensitive to the associated background context.

### 3.3. Univariate Reinstatement Index

The fMRI reinstatement indices were initially subjected to a 2 (age group) x 2 (retrieval task) x 2 (hemisphere) x 2 (ROI) mixed effects ANOVA, the results of which are reported in Table 4. As is evident from the table, none of the main effects reached significance, and likewise, the age group-by-ROI, age group-by-hemisphere, task-by-hemisphere, and ROI-by-task interactions were also not significant. However, the ANOVA did reveal significant interactions between the factors of ROI and hemisphere (of little interest in the current context), and between age group and task. The significant age group-by-task interaction indicated that younger and older adults demonstrated differential scene reinstatement effects according to retrieval task (see Figure 4). Since the factors of hemisphere and ROI did not interact either with age group or retrieval task, reinstatement indices were collapsed across these factors to simplify the follow-up analyses.

**Figure 4.:**
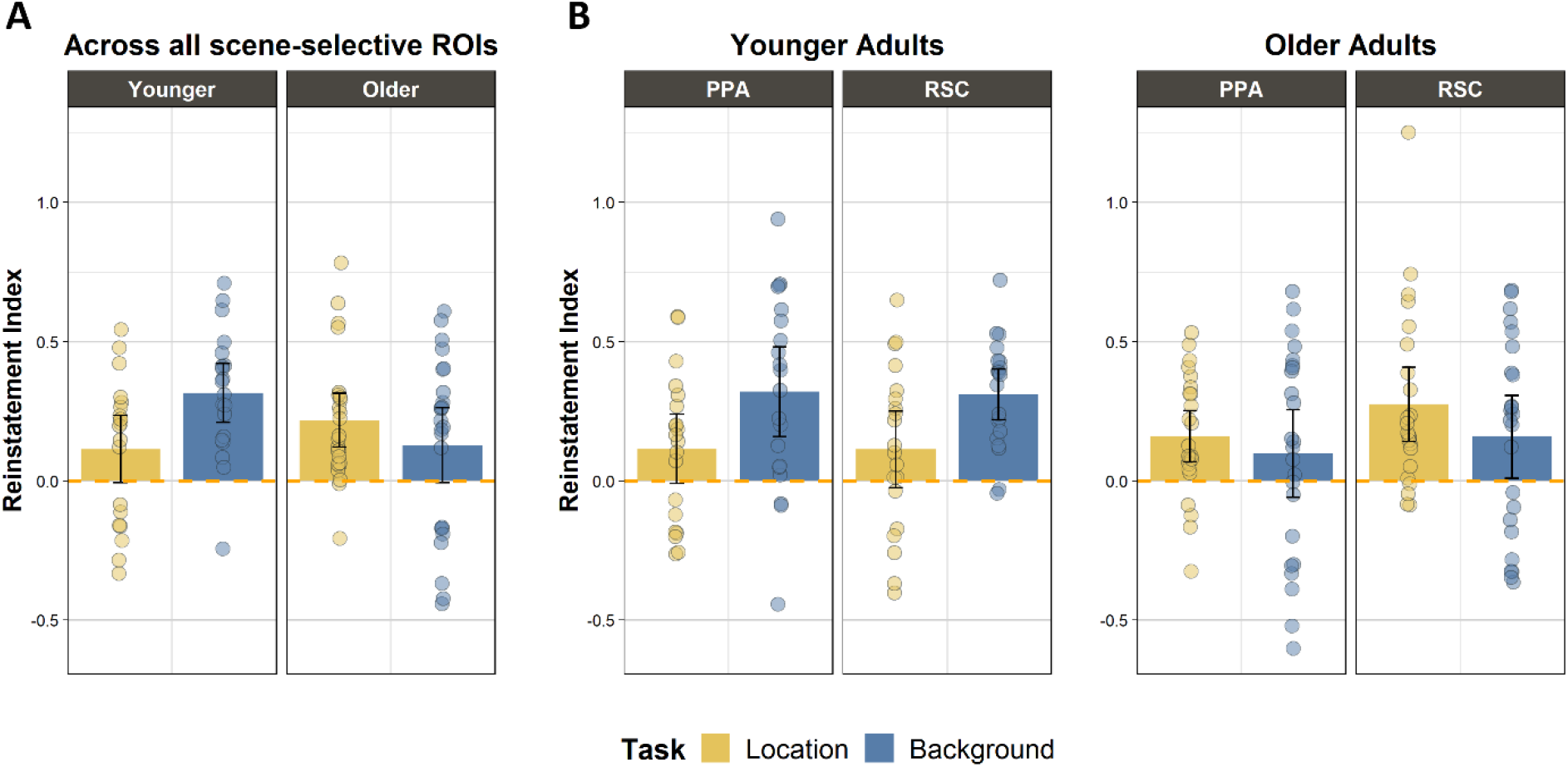
*Reinstatement indices computed separately for the background and location tasks*. **A:** *Reinstatement indices in the two tasks collapsed across ROIs. **B:** Although the main effect of ROI and its interactions with task or age group were not significant, additional panels present the reinstatement indices in the two tasks plotted separately for PPA and RSC for illustrative purposes. In younger adults, task differences were significant in both PPA and RSC (p = 0.015 and p = 0.021, respectively), whereas no task effects were identified in older adults in either of the ROIs (PPA: p = 0.414; RSC: p = 0.101). Error bars represent 95% confidence intervals.*

**Table 4.** Results for the 2 (age group) x 2 (retrieval task) x 2 (hemisphere) x 2 (ROI) mixed ANOVA of the univariate reinstatement index. Significant effect denoted by *.

| | df | F | partial-$\eta^2$ | p-value |
| --- | --- | --- | --- | --- |
| Age group | 1, 42 | 0.405 | 0.010 | 0.528 |
| Task | 1, 42 | 1.542 | 0.035 | 0.221 |
| Hemisphere | 1, 42 | 3.901 | 0.085 | 0.055 |
| ROI | 1, 42 | 1.284 | 0.030 | 0.263 |
| Age group x Task | 1, 42 | 10.496 | 0.200 | 0.002 * |
| Age group x Hemisphere | 1, 42 | 0.719 | 0.017 | 0.401 |
| Age group x ROI | 1, 42 | 1.691 | 0.039 | 0.201 |
| Task x Hemisphere | 1, 42 | 0.395 | 0.009 | 0.533 |
| Task x ROI | 1, 42 | 0.314 | 0.007 | 0.578 |
| Hemisphere x ROI | 1, 42 | 8.832 | 0.174 | 0.005 * |
| Age group x Task x Hemisphere | 1, 42 | 0.064 | 0.002 | 0.802 |
| Age group x Task x ROI | 1, 42 | 0.180 | 0.004 | 0.673 |
| Age group x Hemisphere x ROI | 1, 42 | 1.411 | 0.032 | 0.242 |
| Task x Hemisphere x ROI | 1, 42 | 0.476 | 0.011 | 0.494 |
| Age group x Task x Hemisphere x ROI | 1, 42 | 0.389 | 0.009 | 0.536 |

In light of the results of the foregoing ANOVA we went on to examine scene reinstatement (collapsed across ROIs and hemispheres) as a function of task separately in younger and older adults (Figure 4). In the younger adult group, reinstatement indices were significantly lower in the location relative to the background task (t_(19)_ = 2.973, p = 0.008, d = 0.665). In the older adult group, however, scene reinstatement did not significantly differ as a function of retrieval task (t_(23)_ = 1.508, p = 0.145, d = 0.308), suggesting that reinstatement was not modulated according to the retrieval goal. We evaluated whether reinstatement effects differed reliably from zero as a function of age group and retrieval task with one-sample t-tests (one-tailed). Reliable effects were evident in all cases (Younger Adults: Location task: t_(19)_ = 1.979, p = 0.031, d = 0.443, Background task: t_(19)_ = 6.233, p < 0.001, d = 1.394; Older Adults: Location task: t_(23)_ = 4.661, p < 0.001, d = 0.951, Background task: t_(23)_ = 1.967, p = 0.031, d = 0.402).

### 3.4. Pattern Similarity Analysis

Similarity indices (see PSA Methods) were subjected to a 2 (age group) x 2 (retrieval task) x 2 (hemisphere) x 2 (ROI) mixed ANOVA analogous to that employed for the reinstatement indices. As is indicated in Table 5, none of the four factors gave rise to a significant main effect. Of the interaction effects, only the 2-way interaction between age group and task, and the 3-way interaction between age group, task and ROI reached significance. The age group-by-task interaction parallels the findings for the reinstatement index, in that similarity indices were differentially moderated by retrieval task according to age group (Figure 5-A). Additionally, the significant 3-way interaction between age group, task and ROI indicates that age-dependent task differences differed between the PPA and RSC ROIs.

**Figure 5:**
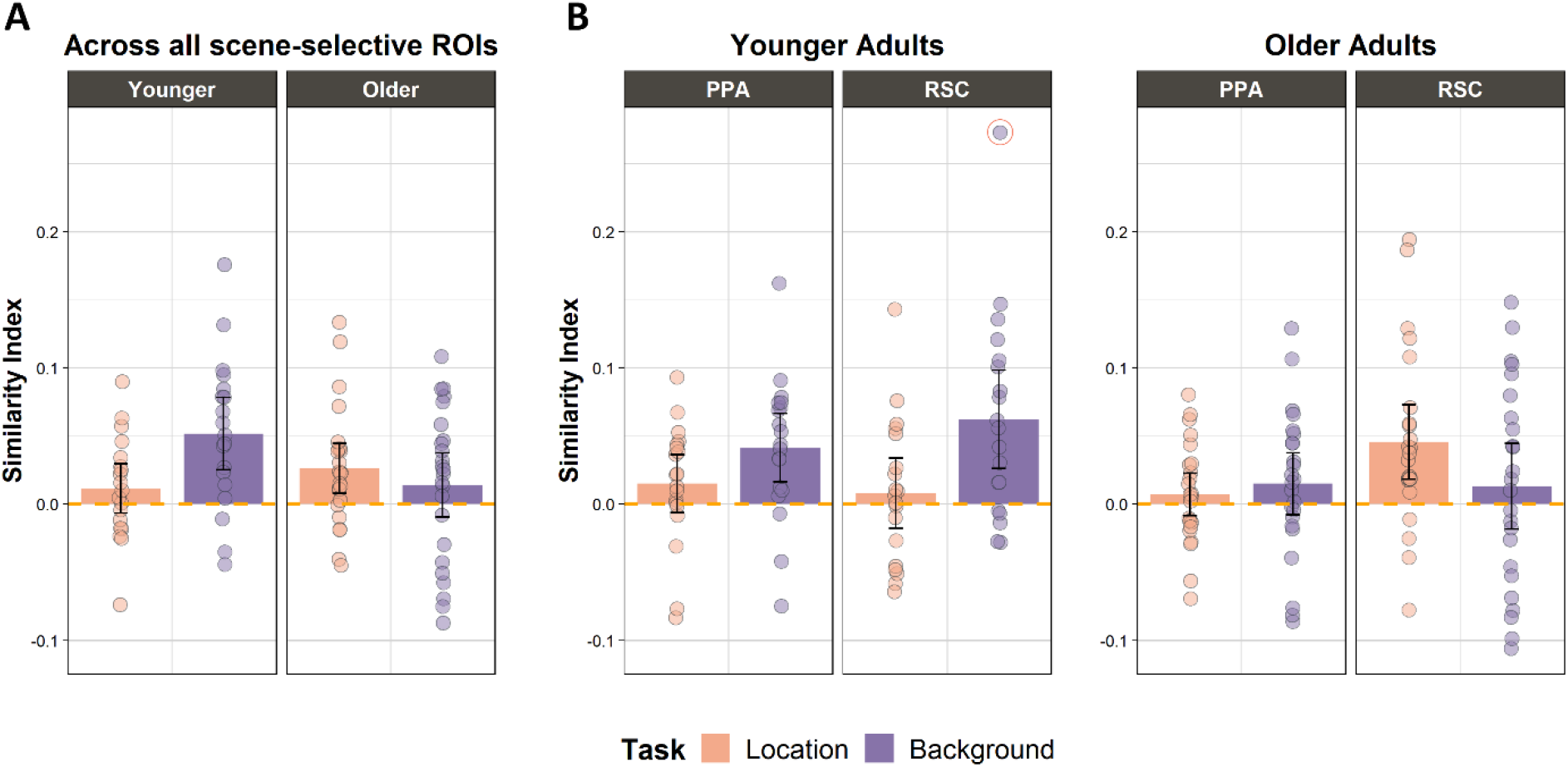
Similarity indices derived from PSA between localizer and memory test. **A:** Similarity index for younger and older adults collapsed across the two ROIs, reflective of the significant age group-by-task interaction. **B:** Similarity index plotted separately for PPA and RSC. Removing the highlighted outlier does not change the outcome of the age group-by-task interaction, nor the main effect of task in the younger cohort. Error bars represent 95% confidence intervals.

**Table 5.** Results for the 2 (age group) x 2 (retrieval task) x 2 (hemisphere) x 2 (ROI) mixed ANOVA of the similarity indices derived from the localizer-test Pattern similarity analysis. Significant effect denoted by *.

| | df | F | partial-$\eta^2$ | p-value |
| --- | --- | --- | --- | --- |
| Age group | 1, 41 | 1.176 | 0.028 | 0.285 |
| Task | 1, 41 | 1.712 | 0.040 | 0.198 |
| Hemisphere | 1, 41 | 0.885 | 0.021 | 0.352 |
| ROI | 1, 41 | 2.679 | 0.061 | 0.109 |
| Age group x Task | 1, 41 | 6.034 | 0.128 | 0.018 * |
| Age group x Hemisphere | 1, 41 | 0.323 | 0.008 | 0.573 |
| Age group x ROI | 1, 41 | 0.553 | 0.013 | 0.461 |
| Task x Hemisphere | 1, 41 | 3.844 | 0.086 | 0.057 |
| Task x ROI | 1, 41 | 0.268 | 0.006 | 0.608 |
| Hemisphere x ROI | 1, 41 | 0.775 | 0.019 | 0.384 |
| Age group x Task x Hemisphere | 1, 41 | 0.127 | 0.003 | 0.723 |
| Age group x Task x ROI | 1, 41 | 8.218 | 0.167 | 0.007 * |
| Age group x Hemisphere x ROI | 1, 41 | 1.169 | 0.028 | 0.286 |
| Task x Hemisphere x ROI | 1, 41 | 0.243 | 0.006 | 0.625 |
| Age group x Task x Hemisphere x ROI | 1, 41 | 1.114 | 0.026 | 0.297 |

Note: One younger adult did not contribute localizer data, and hence was not included in the PSA.

To follow-up the significant 3-way interaction between age group, ROI, and task, additional 2 (ROI) x 2 (retrieval task) ANOVAs were performed separately for the younger and older age groups (Figure 5-B). In the younger adults, the ANOVA identified a significant main effect of retrieval task (F_(1, 18)_ = 5.674, p = 0.028, partial-η^2^ = 0.240), while neither the main effect of ROI (F_(1, 18)_ = 0.368, p = 0.552, partial-η^2^ = 0.020) nor the interaction between task and ROI were significant (F_(1, 18)_ = 2.117, p = 0.163, partial-η^2^ = 0.105). The similarity indices for this group significantly exceeded zero in the background task in both RSC (t_(18)_ = 3.616, p < 0.001, d = 0.830) and the PPA (t_(18)_ = 3.466, p = 0.001, d = 0.795), but did not differ reliably from zero in the location task (PPA: t_(18)_ = 1.488, p = 0.077, d = 0.341; RSC: t_(18)_ = 0.664, p = 0.257, d = 0.152). Thus, the PSA findings for the younger adults complement the results of the reinstatement index and are consistent with the proposal that younger adults gated scene reinstatement during the location task.

In the older adults, the main effects of retrieval task and ROI were not significant (Task: F_(1, 23)_ = 0.822, p = 0.374, partial-η^2^ = 0.035; ROI: F_(1, 23)_ = 3.134, p = 0.090, partial-η^2^ = 0.120). However, the ANOVA revealed a significant interaction between task and ROI (F_(1, 23)_ = 7.461, p = 0.012, partial-η^2^ = 0.245). To unpack this interaction, we performed additional pair-wise contrasts to examine task effects separately in the PPA and RSC. In neither ROI was there a significant difference between the tasks (PPA: t_(23)_ = 0.605, p = 0.551, d = 0.123; RSC: t_(23)_ = 1.844, p = 0.078, d = 0.376). Above-zero similarity indices were identified in the RSC during the location task (t_(23)_ = 3.460, p = 0.001, d = 0.706), whereas the similarity indices in the PPA during the location task, and in both ROIs during the background task, did not differ significantly from zero (ps > 0.093). These findings suggest that although PSA was sensitive to scene reinstatement in older adults, this was confined to the RSC during the location task.

### 3.5. Relationship between scene reinstatement and memory performance

In light of prior research indicating that the strength of cortical reinstatement covaries with the amount and fidelity of retrieved content (e.g. Gordon et al., 2014; Hill et al., 2020; Johnson et al., 2009; Thakral et al., 2015; Trelle et al., 2020), exploratory multiple regression analyses were conducted to examine the relationship between scene reinstatement effects and memory performance. The initial models employed age group, reinstatement indices (separately for each ROI and retrieval task), and the age group-by-reinstatement interaction terms as predictors of memory performance. Because the item memory scores (Pr) for the two tasks were highly correlated (r = 0.844, p < 0.001), and did not significantly differ (see 3.2 Behavioral performance results), we averaged the scores to generate a single estimate of item memory for each participant. In the case of source memory performance, separate regression analyses were conducted for the pSR metrics derived from the respective background and location tasks.

#### 3.5.1. Relationship with item recognition

The regression model predicting Pr from the predictors of age group, PPA reinstatement in the background task, and the interaction term identified a significant interaction between reinstatement and age group (β = 0.806, p < 0.001), indicating that the strength of the relationship between PPA reinstatement during the background task and item memory performance differed between younger and older adults (Figure 6-A). Simple correlations computed for each age group revealed that in younger adults there was a positive relationship between PPA reinstatement and Pr (r = 0.791, p < 0.001), whereas this relationship was absent in older adults (r = - 0.111, p = 0.607; difference between correlations: Z = 3.634, p < 0.001). The remaining 3 regression models (with predictors of age group and the following reinstatement variables: RSC reinstatement in location task, RSC reinstatement in background task, and PPA reinstatement in the location task) did not reveal a significant interaction between age group and reinstatement (ps > 0.086), and reinstatement did not predict Pr when the interaction term was dropped from the models (ps > 0.464).

**Figure 6.:**
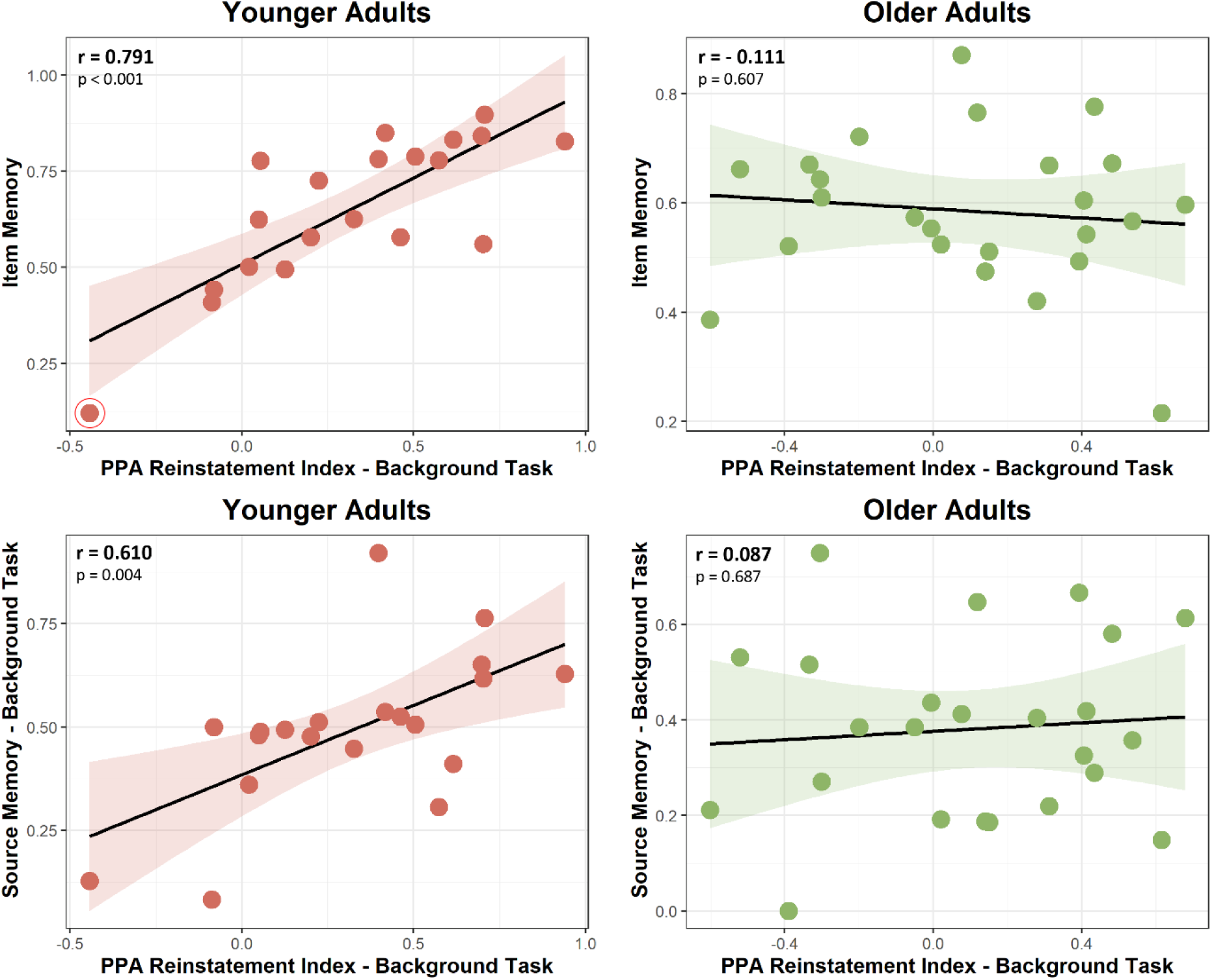
**A**: Scatterplots illustrating zero-order correlations between overall item memory and PPA reinstatement indices derived from trials of the background task. Eliminating the highlighted outlier from the younger adult scatterplot does not change the results (r = 0.698, p < 0.001). **B:** Zero-order correlations between PPA reinstatement of the background task and source memory for backgrounds.

#### 3.5.2. Relationship with source memory

We next performed two regression analyses with age group, reinstatement indices derived during the background task (per ROI), and the interaction term as predictors of background task pSR. Neither PPA nor RSC reinstatement indices for the background task interacted with age group (β = 0.412, p = 0.065, β = 0.125, p = 0.720, respectively). Additionally, RSC reinstatement did not significantly predict background task pSR when the interaction term was omitted (p = 0.345). Because the interaction between age group and PPA reinstatement approached significance, we examined the relationship between PPA reinstatement and pSR in the background task separately for younger and older adults (Figure 6-B). Although the difference between the younger and older adults’ correlations did not quite reach significance (Z = 1.906, p = 0.056), there was a significant positive relationship in the younger (r = 0.610, p = 0.004) but not in the older sample (r = 0.087, p = 0.687).

Moving on to regression models predicting location task pSR, we performed two additional regression analyses employing as predictor variables reinstatement indices in the PPA and RSC derived from the location task. The interaction term was not significant for either ROI (ps > 0.576), and when the interaction term was omitted, the reinstatement indices still did not significantly predict location memory (ps > 0.469). When we repeated the foregoing regression analyses using the similarity indices derived from the PSA as predictors, we were unable to identify any significant relationships with memory performance.

## 4. Discussion

The present study examined potential age differences in a recently identified aspect of retrieval processing termed ‘retrieval gating’ - the ability to regulate the contents of episodic memory retrieval to align them with a retrieval goal. Here, we assessed gating by examining retrieval-related cortical reinstatement of scene information according to whether the information was relevant or irrelevant to the retrieval task. Reinstatement was operationalized with both univariate and multivariate approaches, which converged to suggest that younger, but not older adults, engaged goal-dependent retrieval gating. Additionally, in younger participants only, univariate scene reinstatement indices in the PPA demonstrated strong positive correlations with both scene and item memory. Below, we discuss the significance of these findings for the understanding of the impact of age on controlled episodic retrieval.

### 4.1 Behavioral Results

Turning first to behavioral performance at encoding, we observed no age or background context differences in vividness ratings. Younger adults were however faster to make their vividness responses, and both age groups were faster to make vividness responses to items which were studied over scrambled relative to either of the two scene backgrounds. Under the assumption that the two age groups employed similar criterions when rating vividness of their imagined scenarios, the null effect of age on vividness ratings suggests that differential reinstatement effects in younger and older adults across the two tasks are unlikely to have been confounded by differences in vividness of the imagined scenarios at encoding. With regards to performance at retrieval, item recognition did not differ as a function of age or retrieval task. By contrast, source memory performance was lower in older relative to younger adults. These findings are consistent with prior reports that, while item memory is frequently little affected by age, performance on memory tests that depend heavily on recollection of episodic detail, such as source memory, is consistently impacted (Koen & Yonelinas, 2014; Old & Naveh-Benjamin, 2008; Spencer & Raz, 1995).

Relative to either scene type, test words studied over scrambled backgrounds were associated with better memory for location, but worse memory for the background. These findings are consistent with those reported by Elward & Rugg (2015), who suggested that the lower location memory performance for words studied over scenes was driven by interference from the relatively rich scene information, which could have operated either at encoding or retrieval. According to the first of these possibilities, encoding a word-scene pair is more cognitively demanding than encoding a word presented in association with a scrambled background. The consequent freeing of attentional resources during scrambled trials thus benefited the incidental encoding of the word’s location. This account receives support from the finding that study RTs were shorter for word-scrambled image pairs than for word-scene pairs. Elward and Rugg (2015) also suggested that location memory for words studied over scenes may have suffered because task-irrelevant scene information was insufficiently gated at retrieval and hence interfered with the ability to retrieve or attend to other features of the study episode. The present data arguably speak against this possibility: although the older adults demonstrated little evidence for retrieval gating during the location task (discussed below), the effects of age on source memory performance did not reliably differ between the two retrieval tasks. Under the ‘retrieval interference’ account outlined above older adults should however have been disadvantaged to a greater extent in the location than the background task.

### 4.2. fMRI Results

#### 4.2.1. Age differences in the control of cortical reinstatement

Both younger and older adults demonstrated reliable scene-related cortical reinstatement effects, adding to the already large literature documenting that encoding-related cortical activity is reinstated at retrieval (for reviews see Danker & Anderson, 2010; Rissman & Wagner, 2012; Rugg et al., 2015; Xue, 2018). The present findings in our younger cohort replicate those of Elward & Rugg (2015). Employing an exclusively univariate approach, those researchers reported that younger adults were capable of exerting control over retrieved content, attenuating reinstatement of scene information in the PPA (but not RSC) when it was not task-relevant. The present study replicates the findings observed in the PPA and extends them to the RSC. Additionally, we expand on these findings by demonstrating that the phenomenon of retrieval gating in younger adults extends to multi-voxel indices of reinstatement. The present results join those of Elward and Rugg (2015) in conflicting with the findings reported by Kuhl et al. (2013). Using MVPA classifiers to quantify cortical reinstatement (cf. Johnson et al., 2009), these researchers reported that neither scene nor face reinstatement effects differed according to retrieval goal (which was either to retrieve the face/scene associate of the test word or to retrieve the associate’s study location). The divergence between the results of Kuhl et al. (2013) and the present findings are likely explained by an important procedural difference. At encoding, Kuhl et al. (2013) had their participants view a word presented above two horizontally separated squares, one of which contained an image of a face or a scene. The retrieval phase comprised location and ‘background’ tasks similar to those employed in the present study. Crucially, though, in the location task participants were required to retrieve the location not of the cued word, but of the image that the word had been paired with. Consequently, participants would likely have been motivated to retrieve the image both when prompted to recall its category (face/scene) or its location.

As already noted, unlike younger adults, older adults did not modulate scene reinstatement according to retrieval goal, and hence showed no evidence of retrieval gating (indeed, if anything, there was a non-significant trend towards stronger scene reinstatement in the location task). Thus, the present study provides novel evidence that older adults reinstate features of an episode regardless of whether the information is relevant to the retrieval goal. Although the mechanisms which underlie retrieval gating remain unclear, the present findings are consistent with the notion that the ability to inhibit distracting information declines in older age (Hasher & Zacks, 1988; Hasher et al., 1991; Lustig et al., 2001; see also Campbell et al., 2020). The concept of age deficits in inhibitory control has received much attention in the past, especially given the evidence that older adults’ working memory performance is disproportionately impacted by the presence of distractors (for review see Lustig et al., 2007). This wealth of behavioral studies motivated recent fMRI research which employed working memory paradigms to demonstrate that older adults show less down-regulation of cortical activity in regions responsive to task-irrelevant stimuli (Chadick & Gazzaley, 2011; Chadick et al., 2014; Gazzaley et al., 2005, 2008; Weeks et al., 2020).

It is however yet to be determined whether age differences in retrieval gating reflect a deficit in top-down control or in the ability to implement a feature-selective memory search. It has long been proposed that episodic memory retrieval depends on strategic processes that optimize memory search by targeting goal-relevant information and filtering out irrelevant content (Rugg & Wilding, 2000; Rugg, 2004; Jacoby et al., 2005; see Introduction). From this perspective, the seeming lack of retrieval gating in older adults might reflect an age-related decline in the frontally-mediated control processes that have been proposed to downregulate the representation of intrusive and task-irrelevant information in other experimental settings (e.g. Zanto & Gazzaley, 2019; Zanto et al., 2011). This proposal is consistent with the broader literature linking age-related decline in executive control to reduced structural and functional integrity of prefrontal cortex (Buckner, 2004; Dennis & Cabeza, 2008; Tromp et al., 2015).

#### 4.2.2 Relationship between cortical reinstatement and memory performance

We performed a set of exploratory analyses examining the relationship between cortical reinstatement and memory performance. These analyses were motivated by prior research demonstrating that the strength of reinstatement covaries with the amount of retrieved information (e.g. Gordon et al., 2014; Hill et al., 2020; Johnson et al., 2009; Thakral et al., 2015; Trelle et al., 2020). In the younger cohort, PPA reinstatement in the background task was positively correlated with memory for scenes, as well as with item memory across both retrieval tasks. The reasons for the absence of a relationship between reinstatement and memory performance in the older cohort are currently unclear. We note, however, that prior studies reporting age differences in the ability to adopt specific retrieval orientations (Duverne et al., 2009; Jacoby et al., 2005; Morcom & Rugg, 2004) raise the possibility that the absent relationship between reinstatement and memory performance in the older cohort reflects the employment of different or more variable retrieval strategies during the background task. For example, some older adults might have attempted to retrieve scene information by generating candidate images of scene exemplars when presented with test words that had been paired with a scrambled background. Scene-related activity might also have been elevated on test trials where participants falsely remembered that a word had been studied over a scene rather than a scrambled background (cf. Kurkela & Dennis, 2016). In support of this possibility, we note that in the background task older adults were more likely to endorse a word as having been studied with a target scene when the word had actually been studied with a scrambled background rather than with a non-target scene (probability of incorrect source response [pSource Incorrect] of 0.25 vs. 0.18 respectively; t_(23)_ = 2.277, p = 0.032). This difference was not evident in the younger adult group (pSource incorrect for scrambled = 0.12; pSource incorrect for non-target scene = 0.11; t_(19)_ = 0.639, p = 0.530). Because scrambled trials are the baseline against which reinstatement was assessed, scene-related activity elicited by test words that had not in fact been studied with a scene could have led both to an under-estimation of scene reinstatement effects and a breakdown in the relationship between reinstatement and memory.

As noted above, on the assumption that activity in the PPA scaled with success in retrieving scene information, it is unclear why a relationship between scene reinstatement and memory performance was evident for not only for source memory, but for item memory also. One possibility is that younger adults employed scene context as an additional cue when determining whether a word was previously studied. According to dual-process models, recognition memory is supported by the distinct processes of recollection and familiarity (for reviews see Yonelinas, 2002; Yonelinas et al., 2010) and it is highly likely that, in the present experiment, item memory was supported by both processes. Hence, since the strength of scene reinstatement scales with the likelihood of successful scene recollection, scene reinstatement likely also acted as a proxy for the probability of successful recollection more generally, and hence for the contribution of recollection to item memory. It is worth noting that no relationship between scene reinstatement and memory performance was evident for the location task, consistent with the proposal that scene reinstatement during location trials was downregulated and played little or no role in supporting memory performance on the task.

The foregoing arguments do not elucidate why correlations between scene reinstatement and memory performance were evident for the PPA but not for the RSC. In accounting for this dissociation, we first note that, to our knowledge, it is currently unclear whether scene (or any other) information represented in the RSC is available for the conscious control of behavior. Additionally, or alternatively, the absence of reliable correlations in the RSC might be a consequence of the relatively low fidelity with which scene information appears to be represented in this region; notably, it has been reported that the RSC supports gist-based, ‘schematic’ representations of scenes and spatial contexts, in contrast to the PPA, which supports more fine-grained, detailed representations (Aminoff et al., 2013; Bar, 2004; see also Epstein, 2008). This raises the possibility that, in the present case, the scene representations supported by the RSC were too undifferentiated to permit a determination of whether a reinstated scene should be classified as rural or urban.

### 4.3. Limitations

One important limitation of the present study arises from the cross-sectional nature of the design. Consequently, the findings cannot be unambiguously attributed to the effects of aging rather than to a confounding variable such as a cohort effect (Rugg 2017). We also note the possibility that the correlations identified in the younger adults between PPA scene reinstatement and memory performance - which arguably are unexpectedly high given the inherently noisy nature of neural and behavioral measures – are likely an overestimation of the true effect size (Button et al., 2013). Therefore, these values should be treated with circumspection until a replication study has been reported (Wilson et al., 2020).

The present study does however overcome some of the constraints which commonly apply to studies examining age differences in episodic memory retrieval. Firstly, age differences in retrieval gating cannot be attributed to simple age differences in the strength of cortical reinstatement (Bowman et al., 2019; Folville et al., 2020; Hill et al., 2020; St-Laurent & Buchsbaum, 2019), as both the univariate and multivariate approaches failed to identify a moderating effect of age group on the two metrics of reinstatement (see Thakral et al., 2017; Wang et al., 2016 for similar findings). Secondly, as noted in the methods, the reinstatement index and PSA metrics employed here to quantify reinstatement effects are insensitive to individual differences in HRF gain; hence, the present results are free from the confounding influence of systematic age differences in this variable (e.g. Lu et al., 2011). It should be noted, however, that it remains unclear to what extent the findings might have reflected age differences in the variability or the shape of the hemodynamic response (D’Esposito et al., 2003).

### 4.4. Conclusions

The present study provides novel evidence that younger, but not older adults, can control the contents of recollection in alignment with behavioral goals. Given the novel nature of the concept of retrieval gating, the mechanistic underpinnings of the phenomenon remain to be elucidated. Future research should aim to gain insight into how age-related differences in retrieval gating relate to other processes known to be sensitive to increasing age, such as age deficits in inhibitory control. It also remains unclear whether the failure of the older adults to engage retrieval gating in the present study reflects an *incapacity* to gate as opposed to a preference for a different retrieval strategy. Therefore, future studies should also examine whether older adults can be incentivized to engage retrieval gating strategies (cf. Duverne et al., 2009).

## Acknowledgements

We are grateful to Melanie Racenstein for her assistance with data collection and to our colleagues at the University of Texas Southwestern Medical Center – Advanced Imaging Research Center for their assistance with fMRI data acquisition.

## Author contributions

**Sabina Srokova** – Methodology, Software, Validation, Formal analysis, Investigation, Writing - Original Draft, Review & Editing, Visualization

**Paul F. Hill** – Software, Validation, Writing - Review & Editing, Supervision, Project Administration

**Rachael L. Elward** – Conceptualization, Resources, Writing – Review & Editing

**Michael D. Rugg** – Conceptualization, Methodology, Validation, Writing – Original Draft, Review & Editing, Supervision, Project administration, Funding acquisition

